# Integrated serum proteomics and autoantibody analyses reveal a biomarker signature predictive of flare during biologic tapering in rheumatoid arthritis

**DOI:** 10.64898/2026.05.19.726198

**Authors:** Francisco J. Blanco, Patricia Quaranta, Pablo Domínguez-Guerrero, Valentina Calamia, Patricia Fernández-Puente, Rocío Paz-González, Vanesa Balboa-Barreiro, Diana Noriega, Laura Galindo, Belén Acasuso, Natividad Oreiro, Ricardo Rojo, Lucía Lourido, Cristina Ruiz-Romero

## Abstract

**Background:** Rheumatoid arthritis (RA) is a chronic immune-mediated inflammatory disease characterized by a heterogeneous clinical course with periods of remission and flare. Although biologic DMARDs (bDMARDs) have revolutionized RA treatment by enabling sustained disease control, their long-term use is associated with adverse effects and high costs, making dose tapering an attractive but clinically challenging strategy. The lack of reliable biomarkers to predict flare risk limits safe implementation of treatment de-escalation. This study aimed to identify novel circulating protein biomarkers associated with flare risk in RA patients undergoing bDMARDs tapering, useful to enable biomarker-guided treatment optimization strategies.

**Methods:** A discovery proteomic analysis using mass spectrometry was performed on baseline serum samples from a subset of the OPTIBIO clinical trial (n=44), followed by validation in the full cohort (n=194) using ELISA. Functional pathway analysis explored biological processes associated with candidate biomarkers. In parallel, anti-cytokine autoantibodies were profiled using multiplex immunoassays. Logistic and Cox regression models were used to assess associations with flare risk. Predictive models integrating biomarkers and clinical variables were evaluated using receiver operating characteristic (ROC) analysis, sensitivity and specificity metrics, and decision curve analysis to assess clinical utility.

**Results:** Mass spectrometry identified 806 proteins, of which 87 were differentially expressed at baseline between patients who flared and those who maintained remission during follow-up within the intervention (tapering) arm. Functional enrichment analysis highlighted immune-regulatory and innate immune pathways. Among the candidates, V-set immunoglobulin-domain-containing 4 (VSIG4) was validated as a biomarker associated with increased flare risk. Anti-interferon-γ (anti-IFNγ) autoantibodies were also associated with flare. A combined model including VSIG4, anti-IFNγ, and the clinical variable DAS28-CRP improved predictive performance compared with clinical variables alone (AUC 0.76 vs 0.66), achieving significantly higher sensitivity. Decision curve analysis demonstrated higher net benefit of the combined model, indicating improved clinical decision-making. In a secondary analysis focused on patients with prolonged remission, representing the most suitable candidates for safe treatment tapering, the model performance further improved (AUC 0.84).

**Conclusion:** Integration of novel serum proteomic and autoantibody biomarkers with clinical parameters improves prediction of flare during biologic tapering in RA and provides clinically relevant benefit for patient stratification. These findings support further development of biomarker-driven approaches for personalized treatment optimization strategies.

## BACKGROUND

Rheumatoid arthritis (RA) is a chronic autoimmune disorder characterized by inflammation and hyperplasia of the synovial membrane, autoantibody production, and progressive damage to cartilage and bone [1]. It can also present with systemic manifestations affecting the cardiovascular and respiratory systems. The course of RA is often marked by fluctuations in disease activity, including episodes of sudden worsening or reactivation of underlying inflammatory mechanisms, known as flares, which can have a wide range of adverse effects on patients [2]. These flares are associated with reduced physical function, increased fatigue, and impaired quality of life, as well as long-term consequences such as progressive joint damage and an increased risk of cardiovascular disease [3].

Despite their clinical importance, the biological basis of RA flares remains poorly understood, and their unpredictable and sporadic nature makes them difficult to investigate [4]. Historically, frequent flares were common among patients with RA [1]. Today, early diagnosis combined with timely treatment using biological disease-modifying anti-rheumatic drugs (bDMARDs) enables approximately half of patients to achieve sustained remission [5]. However, bDMARDs are associated with potential risks, high cost, and the need of continuous monitoring through regular blood tests [1]. Consequently, clinical guidelines include recommendations for tapering bDMARDs in RA. In Europe, dose reduction may be considered after at least six months of glucocorticoid-free remission, whereas in the United States maintenance of the therapeutic dose is generally recommended due to limited evidence supporting the safety and efficacy of tapering strategies [6,7]. Within this context, the OPTIBIO study (Eudra-CT 2012-004482-40) was a phase IV, randomized, controlled, open-label, non-inferiority trial conducted in patients with RA in sustained remission on biologic therapy. The study aimed to compare flare rates at 12 months and to identify predictors of successful treatment optimization. The results showed that bDMARD dose reduction was not non-inferior to continued therapy, although both strategies demonstrated comparable safety profiles [8].

Although tapering is desirable, reliable tools to identify patients suitable for safe tapering while minimizing relapse risk are still lacking [9]. The identification of predictors of flare during bDMARD tapering has proven challenging. Known risk factors include seropositivity, shared epitope, high baseline disease activity, and longer disease duration [10]. Subclinical inflammation is also considered an important predictor, but proposed biomarkers such as multibiomarker scores and imaging approaches have shown inconsistent performance and lack validation [4,11,12].

Given the substantial clinical and socioeconomic impact of bDMARD therapy, identifying biomarkers predictive of relapse is essential to optimize RA management [13]. In this study, we applied a proteomic approach to the OPTIBIO cohort to identify circulating proteins and anti-cytokine autoantibodies associated with flare risk during bDMARD tapering. Conducting this analysis within a controlled trial provides a unique opportunity to gain insight into the immunological mechanisms underlying disease flares and to support the development of biomarker-guided strategies for personalized treatment optimization.

## METHODS

### Study population

All procedures were conducted according to the Declaration of Helsinki. Written informed consent was obtained from all participants before inclusion. In this study, we analyzed baseline serum samples from 194 patients with RA enrolled in the OPTIBIO study (Eudra-CT 2012-004482-40) [8]. Baseline characteristics of the patients are summarized in **Table 1**. This trial was designed to assess whether reduced-dose biological therapy could maintain remission in RA patients and was conducted in five Spanish hospitals. Adults in remission for ≥6 months (DAS28 <2.6, SDAI <5, or ACR/EULAR 2011 criteria) were treated with TNF inhibitors, Tocilizumab, or Abatacept. Participants were randomly assigned either to an intervention group with dose tapering or to a control group receiving standard therapy. Flares were defined as DAS28 >2.6, SDAI >5, or failure to meet remission criteria. Patients were followed for 1–3 years, with assessments every 12 weeks. The trial evaluated both remission maintenance and the non-inferiority of reduced-dose regimens.

**Table 1.**
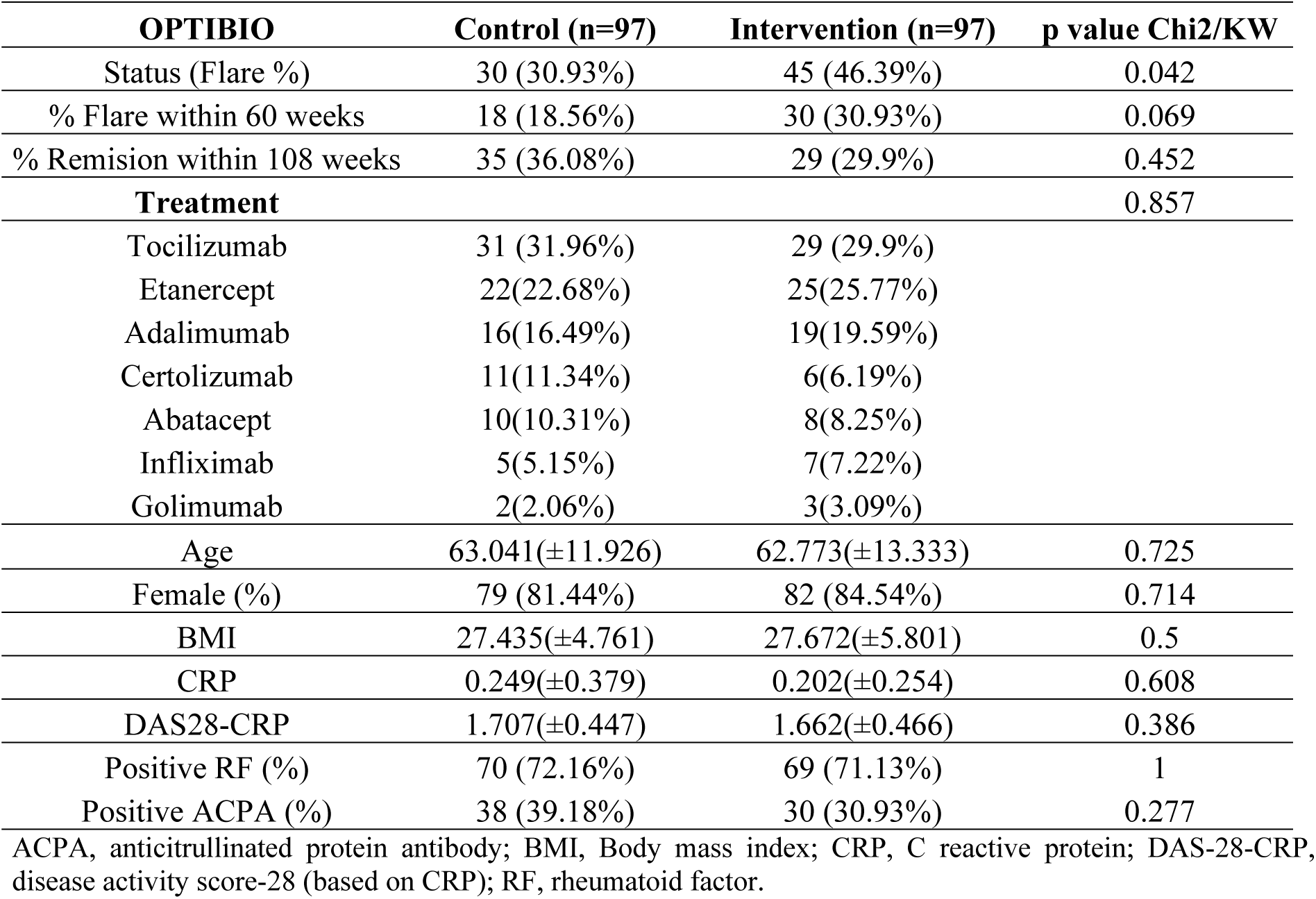
Baseline characteristics of patients from the OPTIBIO cohort analyzed in this study.

| OPTIBIO | Control (n=97) | Intervention (n=97) | p value Chi2/KW |
| --- | --- | --- | --- |
| Status (Flare %) | 30 (30.93%) | 45 (46.39%) | 0.042 |
| % Flare within 60 weeks | 18 (18.56%) | 30 (30.93%) | 0.069 |
| % Remission within 108 weeks | 35 (36.08%) | 29 (29.9%) | 0.452 |
| **Treatment** |  |  | 0.857 |
| Tocilizumab | 31 (31.96%) | 29 (29.9%) |  |
| Etanercept | 22(22.68%) | 25(25.77%) |  |
| Adalimumab | 16(16.49%) | 19(19.59%) |  |
| Certolizumab | 11(11.34%) | 6(6.19%) |  |
| Abatacept | 10(10.31%) | 8(8.25%) |  |
| Infliximab | 5(5.15%) | 7(7.22%) |  |
| Golimumab | 2(2.06%) | 3(3.09%) |  |
| Age | 63.041(±11.926) | 62.773(±13.333) | 0.725 |
| Female (%) | 79 (81.44%) | 82 (84.54%) | 0.714 |
| BMI | 27.435(±4.761) | 27.672(±5.801) | 0.5 |
| CRP | 0.249(±0.379) | 0.202(±0.254) | 0.608 |
| DAS28-CRP | 1.707(±0.447) | 1.662(±0.466) | 0.386 |
| Positive RF (%) | 70 (72.16%) | 69 (71.13%) | 1 |
| Positive ACPA (%) | 38 (39.18%) | 30 (30.93%) | 0.277 |

ACPA, anticitrullinated protein antibody; BMI, Body mass index; CRP, C reactive protein; DAS-28-CRP, disease activity score-28 (based on CRP); RF, rheumatoid factor.

### Sample processing for mass spectrometry (MS)

Of the 194 serum samples, 44 were selected for the discovery phase of biomarker identification using MS, ensuring balance in flare status, treatment group, and ACPA positivity (**Supplementary Table 1**). After thawing and centrifugation, the most abundant serum proteins were depleted using High Select Top14 Abundant Protein Depletion spin columns (Thermo Fisher, MA, USA) to improve sensitivity and proteome coverage. Protein concentrations were measured with NanoDrop at 280 nm, and 10 μg of protein per sample underwent in-solution digestion. Briefly, proteins were denatured, reduced with DTT, alkylated with iodoacetamide, diluted, and digested overnight with trypsin. The resulting peptides were acidified, desalted using in-house C18 stage tips, and dried.

Proteomic profiling was performed by label-free shotgun analysis on a timsTOF Pro mass spectrometer coupled to a nanoElute LC system (TIMS TOF Pro LC-MS/MS System, Bruker, MA, USA). Peptides were separated by reversed-phase chromatography with a linear gradient and analysed using data-independent acquisition (DIA) with PASEF. Quality control included blank runs between samples and technical triplicates of one randomly selected sample. The mass spectrometry proteomics raw data have been deposited to the ProteomeXchange Consortium via the PRIDE [14] partner repository with the dataset identifier PXD073940.

Quantitative analysis of the 44 DIA runs was carried out using Spectronaut™, searching against the UniProtKB/Swiss-Prot database with default settings [15]. Protein identification required a false discovery rate (FDR) <0.01. Significant changes were defined as q-value <0.05 with log2 fold change >0.58 or <-0.58. Differentially expressed proteins were further investigated through functional enrichment analysis using STRING.

### ELISA measurements

Circulating calprotectin (S100A8/S100A9 heterodimer) and V-set immunoglobulin-domain-containing 4 (VSIG4) were measured in the 194 serum samples using commercially available ELISA kits. Calprotectin was measured with the Human S100A8/S100A9 Heterodimer ELISA Kit – Quantikine purchased from R&D systems (Minneapolis, MN, USA), and VSIG4 measured with an ELISA kit from RayBiotech life (Peachtree Corners, GA, USA). Signals were read using an Infinite M200 Nanoquant plate reader (Tecan) at 450nm using 570nm as a reference. Four-parameter logistic (4PL) standard curves were generated from serial dilutions of standards to calculate serum protein concentrations.

### Protein microarray analysis

A bead-based antigen array (MILLIPLEX® MAP Human Cytokine Autoantibody Magnetic Bead Panel - HCYTAAB-17K, Merck, Darmstadt, Germany) was used for the simultaneous detection and semi-quantification of 15 anti-cytokine autoantibodies (ACAAs) in the 194 serum samples of the OPTIBIO cohort: anti-BAFF, anti-G-CSF, anti-IFNβ, anti-IFNγ, anti-IL-1α, anti-IL-6, anti-IL-8, anti-IL-10, anti-IL-12 (p40), anti-IL-15, anti-IL-17A, anti-IL-17F, anti-IL-18, anti-IL-22, and anti-TNFα. This array was selected because it includes cytokines known to play key roles in immune regulation and inflammation in RA.

The suspension bead array was prepared according to the manufacturer’s instructions. Briefly, serum samples were thawed, vortexed, centrifuged, and diluted 1:100 in assay buffer. Diluted samples were randomized in 96-welll plates and incubated overnight at 4°C with the bead array by shaking. After incubation, beads were washed, and the antigen-antibody interaction was detected by incubation with an anti-human IgG Fc antibody labeled with the fluorophore phycoerythrin (PE). Finally, beads were washed again, resuspended in Drive Fluid and analyzed using the MagPix instrument (MAGPIX^®^ System, Luminex, TX, USA). Fluorescence signals, corresponding ACAAs, were reported per bead identity as median fluorescence intensities (MFI), reflecting semi-quantitative autoantibody levels in each sample. Assay precision was assessed using coefficients of variation across replicates.

### Statistical analysis

Data were analyzed and visualized using SPSS and R software (version 4.5.1, R Foundation for Statistical Computing). Serum levels of ACAAs (MFI), calprotectin (ng/mL), and VSIG4 (ng/mL), obtained by protein microarrays and ELISA, were compared between the groups using non-parametric tests. Univariable binary logistic regression was used to assess associations between candidate biomarkers and disease flare during follow-up.

The Kaplan Meier method was used to estimate the cumulative probability of flare after tapering (survival function) according to VSIG4 and anti-IFNγ levels. Both biomarkers were categorized into quartiles (VSIG4) or tertiles (anti-IFNγ), with the lowest levels at baseline serving as reference. Intermediate and high categories were compared with the reference group. A multivariable extended Cox proportional hazards model was also fitted, and 95% confidence intervals (CI) for hazard ratios (HR) were reported.

Additionally, univariate logistic regression was performed to define a clinical model to predict flare in the subgroup where anti-IFNγ and VSIG4 showed the highest discriminatory capacity: patients who flared within 60 weeks (1 year) and those who maintained remission beyond 108 weeks (2 years) after tapering. Odds Ratios (OR) with their corresponding 95% confidence interval (CI) were calculated for VSIG4 and Anti-IFNγ. The OR were calculated per 10 ng/mL for VSIG4 absolute quantification data, and per 1000 MFI units in the case of Anti- IFNγ. Predictive performance of the clinical, biomarker, and combined models was evaluated using receiver operating characteristic (ROC) curve analysis. Area under the curve (AUC), sensitivity, specificity, positive predictive value (PPV), negative predictive value (NPV), and accuracy were calculated with corresponding 95% confidence intervals. Optimal probability thresholds for model classification were selected using the Youden index. Model comparison was performed using the DeLong method [16]. Internal validation was performed using bootstrap resampling (B = 500 replicates). Bootstrap-corrected AUC values were calculated with the pROC and boot packages in R. In addition, decision curve analysis (DCA) and net interventions avoided (NIA) were used to assess the potential clinical utility of the models across clinically relevant threshold probabilities. To ensure robust model development, the TRIPOD (Transparent Reporting of a multivariate prediction model for Individual Prognosis Or Diagnosis) guidelines were followed [17].

## RESULTS

### Proteomic discovery of circulating biomarkers associated with flare risk

A proteomic discovery phase using MS was carried out on a subset of 44 serum samples collected at baseline from the OPTIBIO cohort (control group, n=22; intervention group, n=22). The clinical and demographic characteristics of this subgroup of patients is shown in Supplementary Table 1. This analysis identified a total of 806 proteins with at least one unique tryptic peptide (Supplementary Table 2). The coefficient of variance (CV) for the pooled sample injected in triplicate across different runs was 0.25%, demonstrating the reproducibility of the assay. Of the 806 proteins identified, 789 were quantified within the control arm (Supplementary Table 3), and 759 in the intervention arm (Supplementary Table 4). From these, 87 proteins differed significantly between patients who flared (F) and those who remained in remission (R) exclusively in the intervention group, with log2 fold change >0.58 or <-0.58, Q < 0.1 (Figure 1A). These proteins were thus identified as candidate biomarkers associated with flare specifically in bDMARD tapering strategies.

**Figure 1.**
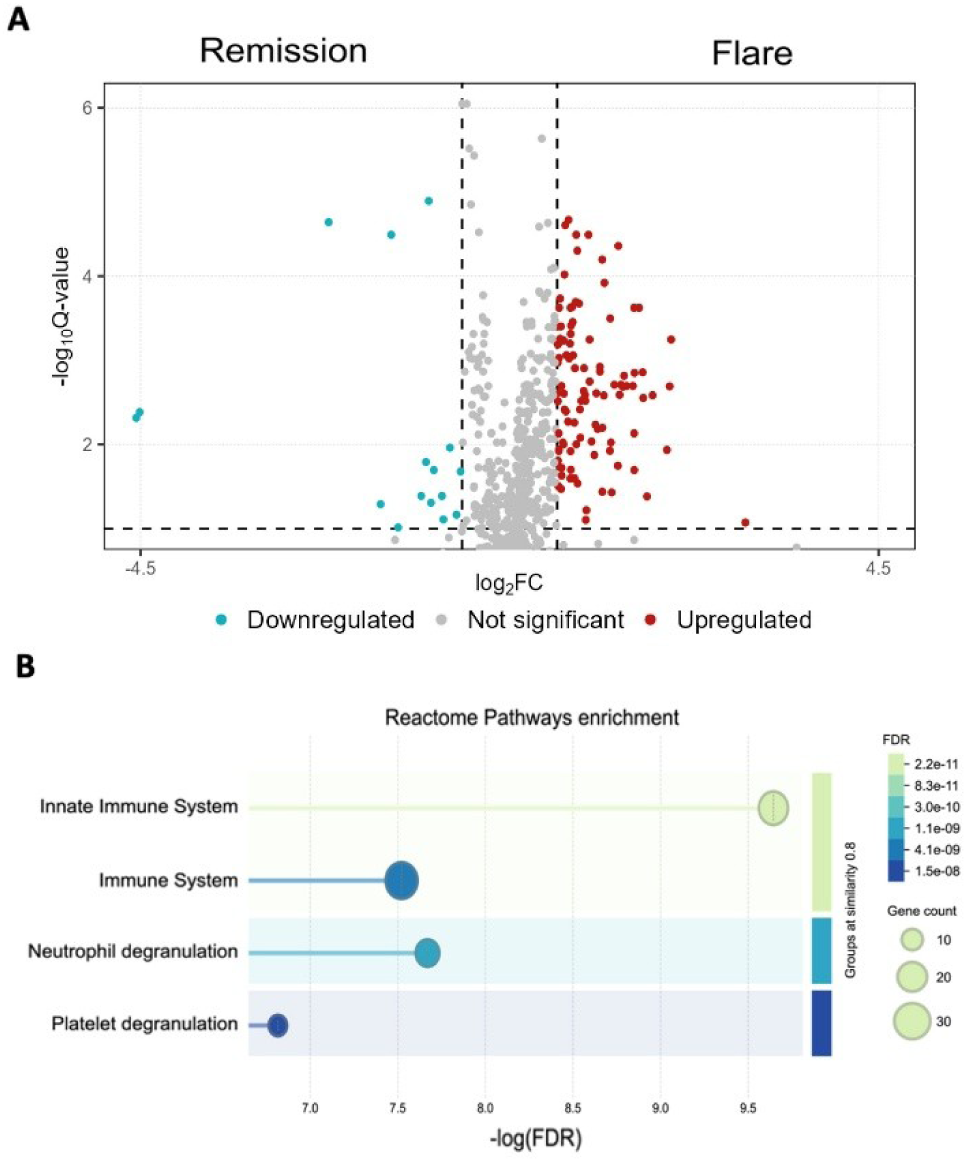
Proteomic discovery of predictive biomarkers of flare in DMARD tapering strategies. (A) Volcano plot showing upregulated (red) and downregulated (blue) proteins identified by MS at baseline in patients who flared (n=11) *versus* those who stayed in remission (n=11) in the intervention arm. Cutoffs were set at log2 fold change ≥0.58 and q-value < 0.1. (B) Biological process enrichment of quantified and differentially modulated proteins (q-value <0.1). The Y-axis shows enriched biological processes, and the X-axis represents the proportion of proteins associated with each process relative to all proteins annotated in that category. Bubble size reflects the number of proteins involved, and bubble color indicates the false discovery rate (FDR), with lower values representing higher statistical significance.

Reactome pathway enrichment analysis on this subset of 87 proteins revealed a strong overrepresentation of immune-related pathways (Figure 1B). As illustrated, around 35 proteins were associated with immune system functions, with the innate immune response displaying the lowest false discovery rate (FDR). Additionally, processes such as neutrophil and platelet degranulation, which contribute to inflammation, were also highlighted in the enrichment analysis.

### Validation of VSIG4 as a protein biomarker associated with flare risk

To validate findings from the discovery phase, VSIG4 (V-set immunoglobulin-domain-containing 4) and the heterodimer S100A8/S100A9 (calprotectin) were analyzed by ELISA in the full OPTIBIO cohort (n=194, characteristics shown in Table 1). These proteins were prioritized for validation based on their reported roles in RA pathophysiology [18,19], supporting their mechanistic relevance, together with their modulation observed within the intervention arm in the discovery analysis.

Median baseline VSIG4 levels were similar between control and intervention groups (73.375 ± 44.440 vs. 74.068± 47.107 ng/mL, respectively). No significant differences were observed between F and R patients within the control group (n = 97) (Figure 2A). However, in the intervention group (n = 97), baseline VSIG4 levels were significantly higher in F patients compared with R (p = 0.017), thus supporting the findings from the discovery phase. When categorized according to the median, higher baseline VSIG4 levels were significantly associated with an increased risk of flare. Logistic regression analysis showed that patients in the higher VSIG4 group had significantly increased odds of flare [OR = 5.07 (95% CI: 2.13–12.06), p < 0.001]. Similarly, elevated VSIG4 levels were associated with a higher risk of flare over time [HR = 3.25 (95% CI: 1.70–6.24), p < 0.001], and Kaplan–Meier analysis demonstrated significantly reduced flare-free survival in patients with high baseline VSIG4 levels compared with those with low levels (log-rank p = 0.000096), indicating that patients undergoing tapering with elevated baseline VSIG4 levels experienced earlier flares during follow-up (Figure 2B).

**Figure 2.**
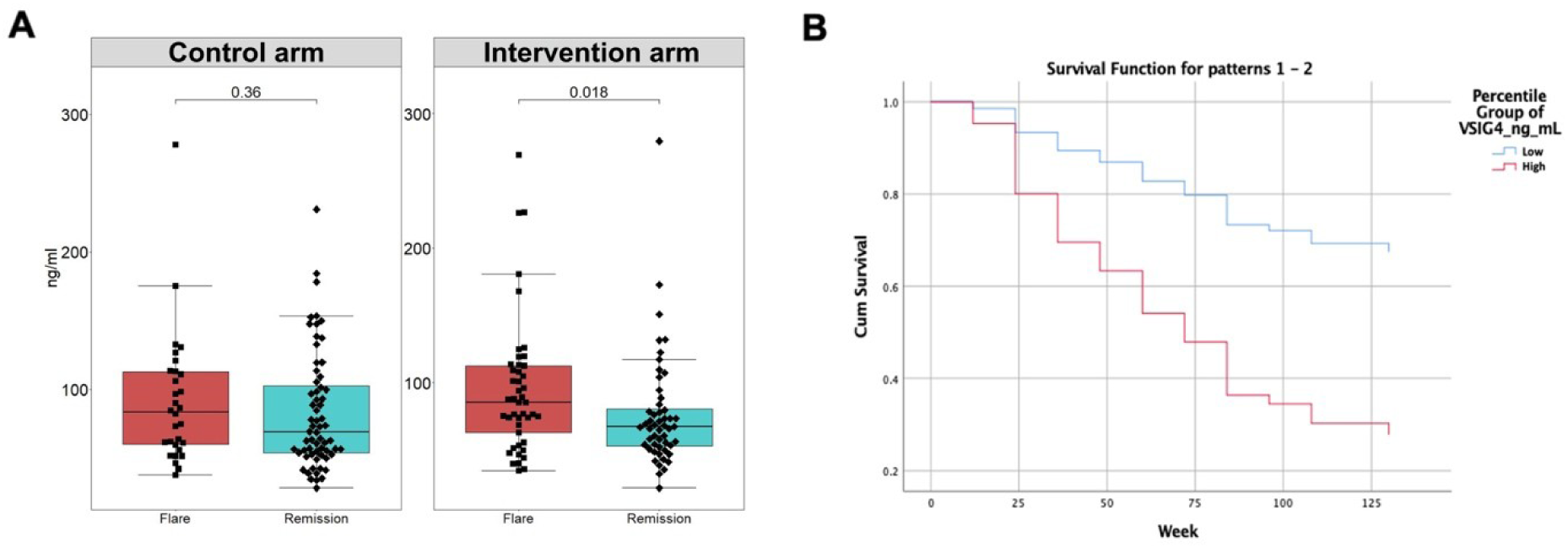
Association of V-set Immunoglobulin-domain-containing 4 (VSIG4) with flare after DMARD tapering. Circulating VSIG4 levels (ng/mL) at the time of DMARD tapering, stratified by remission (blue) and flare (red) occurrence in patients in the control arm (left), and patients in the intervention arm (right). P-values were calculated using Mann-Whitney U tests. Bars represent median and interquartile range. (B) Kaplan-Meier curve stratified by baseline VSIG4 levels (low, blue; high, red), showing a significantly reduced flare-free survival in patients with high VSIG4 levels undergoing tapering.

In contrast, baseline calprotectin levels showed no significant differences between F and R patients in both the control and intervention groups of the trial (Supplementary Figure 1A-B). Median calprotectin levels were also comparable between groups (6963.68 ± 3221.23 in the control group, 6828.52 ± 4036.36 ng/mL in the intervention group), with no significant differences observed.

### Screening of anti-cytokine autoantibodies associated with flare risk

In parallel, to further investigate the potential contribution of immune-regulatory factors to flare risk, anti-cytokine autoantibodies (ACAAs) were analyzed in baseline serum samples from the full OPTIBIO cohort (n=194). IgG autoantibodies against 15 cytokines known to play key roles in immune regulation and inflammation in RA were profiled using a multiplex antigen array designed for their indirect semi-quantification (see Methods).

Non-parametric analyses showed no significant differences in median ACAA levels between the control and intervention groups (Supplementary Figure 2). However, within the intervention group, anti-IFNγ levels were modestly higher in F patients compared to R (p=0.07) (Figure 3A). In this tapering group, anti-IFNγ levels (scaled per 1000 MFI units) were significantly associated with an increased flare risk during follow-up [OR=1.889 (95% CI: 1.888-1.890), p = 0.019]. When anti-IFNγ levels were categorized into tertiles, patients in the highest tertile had a significantly greater risk of flare over time compared with those in the lowest tertile [HR=2.114 (95% CI: 1.020-4.379), p = 0.044], indicating an earlier occurrence of flare in patients with higher baseline anti-IFNγ levels. A significantly reduced flare-free survival in patients with high baseline anti-IFNγ levels compared with those with low levels (log-rank p = 0.021), whereas no significant differences were observed between intermediate and low anti-IFNγ levels (Figure 3B).

**Figure 3.**
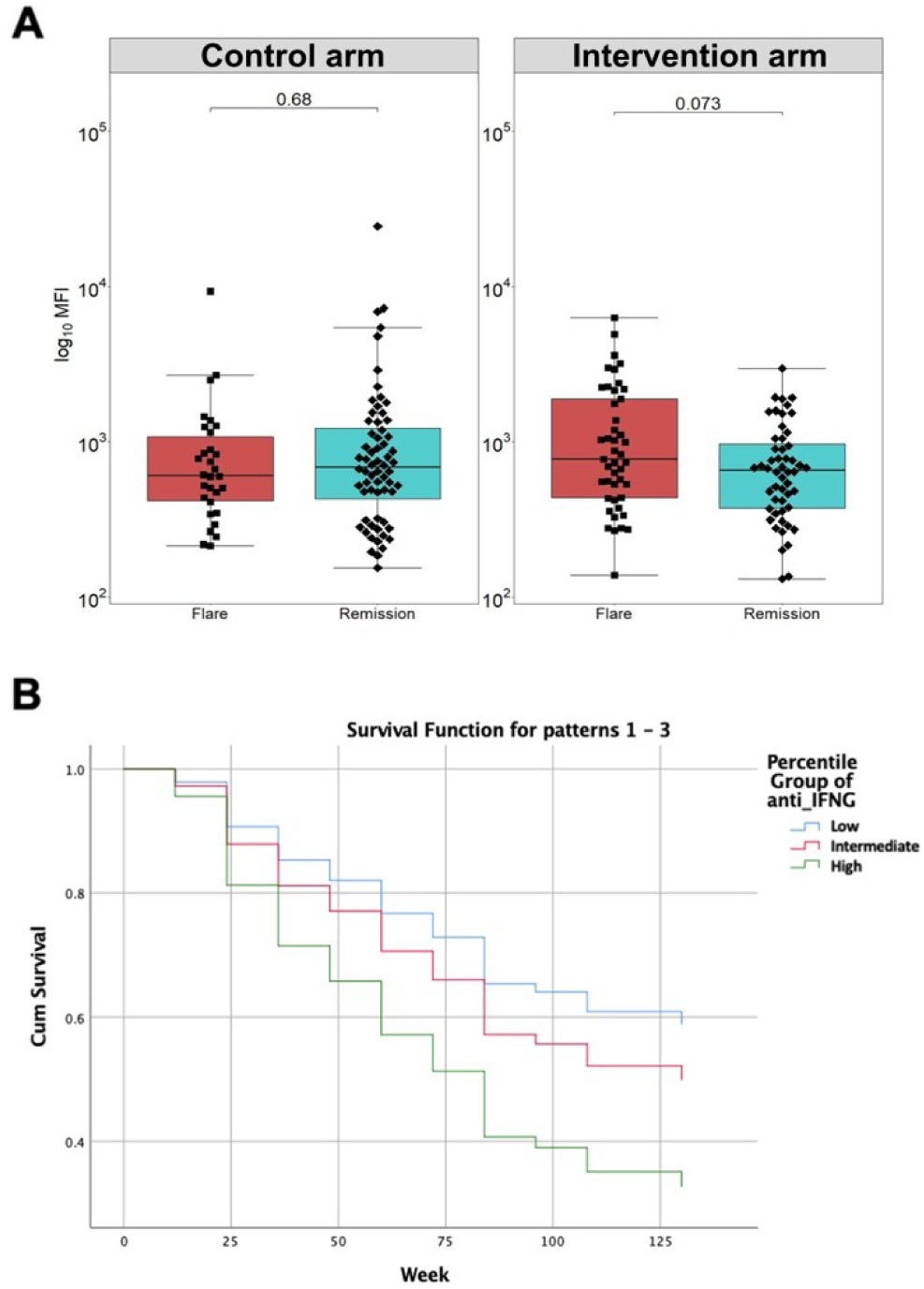
Association of IgG anti-IFNγ levels with flare risk after DMARD tapering. **(A)** Circulating IgG anti-IFNγ at the time of DMARD tapering, stratified by remission (blue) and flare (red) occurrence in patients in the control arm (left), and patients in the intervention arm (right). P-values were calculated using Mann-Whitney U tests. Bars represent median and interquartile range. (B) Kaplan-Meier curve showing flare probability over time in the intervention arm, stratified according to baseline anti-IFNγ levels (low, blue; intermediate, red; high, green). Patients with high anti-IFNγ levels showed significantly reduced flare-free survival compared with those with low levels.

### Development of integrated biomarker and clinical models for flare risk

A univariate logistic regression analysis was first performed to assess the association of demographic and clinical covariates with flare risk in the intervention arm (Supplementary Table 5). Among the variables analyzed, both BMI and baseline DAS28-CRP reached statistical significance (p < 0.05). However, Pearson correlation analysis revealed a significant association between these variables (p = 0.04). In addition, BMI lost statistical significance when included in multivariable models, whereas DAS28-CRP remained independently associated with flare risk. Therefore, only DAS28-CRP was retained in the final clinical model.

Based on these findings, three predictive models were constructed: a clinical model including DAS28-CRP, a biomarker model including VSIG4 and anti-IFNγ, and a combined model integrating both clinical and biomarker variables. Detailed performance metrics of the models are summarized in Supplementary Table 6. In the intervention (tapering) arm, the clinical model including DAS28-CRP was significantly associated with flare risk [OR = 4.44 (95% CI: 1.71–13.17), p = 0.004], yielding an AUC of 0.666 (95% CI: 0.557-0.774). The biomarker model including VSIG4 and anti-IFNγ improved discriminative performance [AUC = 0.722 (95% CI: 0.62-0.823)], while integration of these biomarkers with DAS28-CRP into a combined model further increased predictive accuracy, resulting in an AUC of 0.760 (95% CI: 0.666-0.854) (Figure 4A).

**Figure 4.**
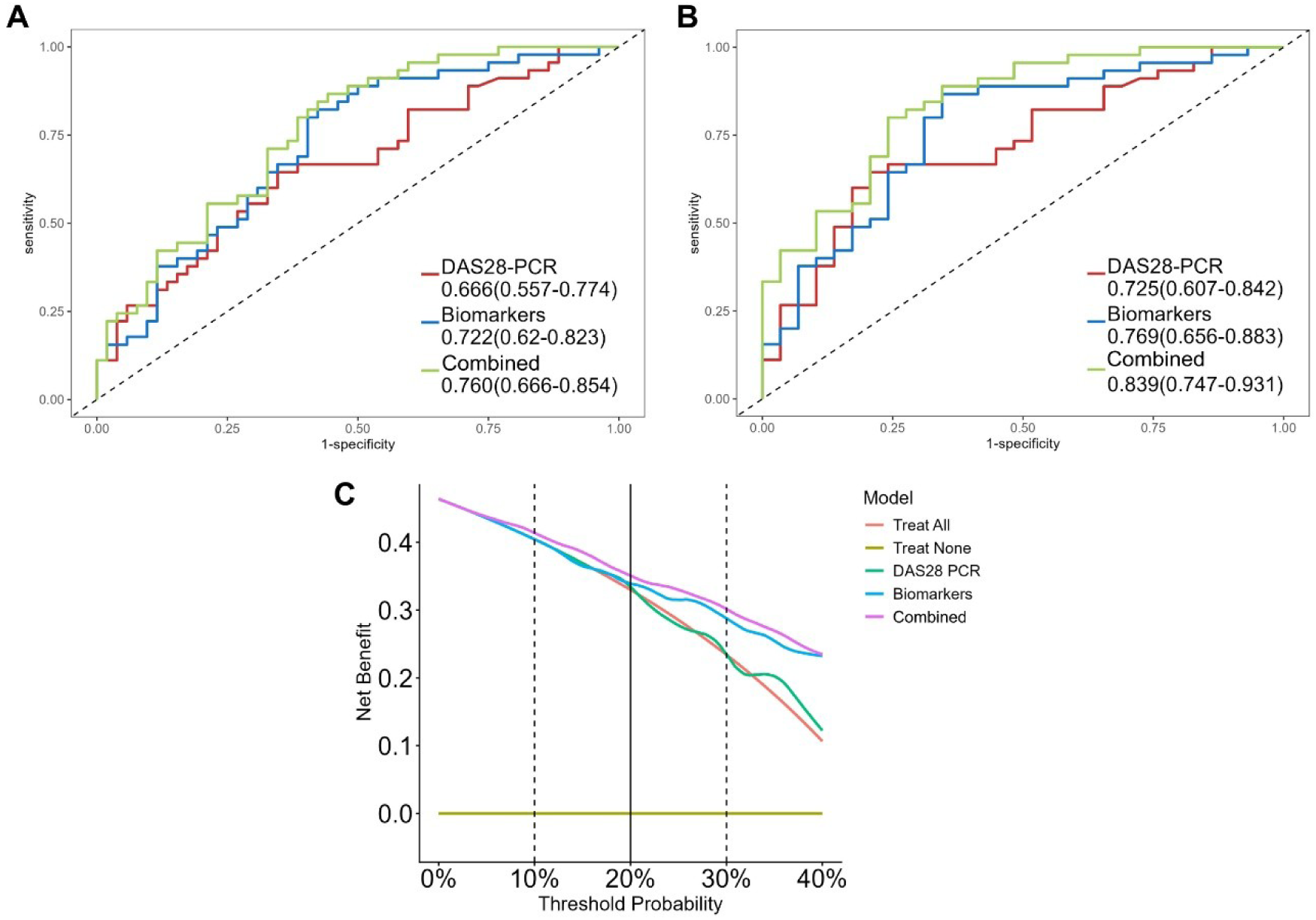
Predictive performance and clinical utility of biomarker-based models for flare risk during bDMARD tapering in RA patients. (A) Receiver operating characteristic (ROC) curves comparing the discriminative performance of the clinical model based on DAS28-CRP, the biomarker model including VSIG4 and anti-IFNγ, and the combined model integrating clinical and biomarker variables in the entire intervention arm of the OPTIBIO cohort. The combined model showed the highest predictive performance (AUC = 0.760). (B) ROC curves for the secondary analysis focused on patients with prolonged remission, representing those most likely to benefit from treatment optimization. In this subgroup, the combined model achieved improved discriminative capacity (AUC = 0.839). (C) Decision curve analysis (DCA) performed in the intervention arm, evaluating the clinical utility of the predictive models across clinically relevant threshold probabilities ranging from 0% to 40%. The combined model demonstrated the highest net benefit compared with the clinical model, the biomarkers model, and the “treat all” and “treat none” strategies.

To further refine patient stratification, a secondary analysis was performed comparing patients who experienced flare with those who maintained prolonged remission (≥2 years), who would be the best candidates for tapering approaches. In this setting (n = 74), all variables showed stronger associations, and the combined model demonstrated markedly improved performance, with VSIG4 [OR = 3.98 (95% CI: 1.26–13.78), p = 0.022], anti-IFNγ [OR = 5.07 (95% CI: 5.06–5.07), p = 0.006], and DAS28-CRP [OR = 10.51 (95% CI: 2.39–65.88), p = 0.005]. This combined model achieved a high discriminative capacity (AUC = 0.839, 95% CI: 0.747-0.931; bootstrapped-adjusted AUC = 0.815, 95% CI: 0.735-0.91) (Figure 4B and Supplementary Table 6). Overall, these results indicate that the integration of serum proteomic biomarkers with clinical parameters improves discrimination of flare risk and enhances the identification of patients who maintain prolonged remission under tapering conditions.

Model performance was further assessed using sensitivity and specificity at the selected probability thresholds determined by the Youden index (Table 2). The biomarker-only model demonstrated high sensitivity (0.822, 95% CI: 0.679-0.920), indicating a strong ability to identify patients who subsequently flared, although with moderate specificity (0.577, 95% CI: 0.432-0.713). The clinical model based on DAS28-CRP showed lower sensitivity (0.644, 96% CI: 0.488-0.781) with similar specificity (0.654, 95% CI: 0.509-0.780). Notably, the combined model integrating VSIG4, anti-IFNγ, and DAS28-CRP achieved the highest sensitivity (0.867, 95% CI: 0.732-0.949) while maintaining acceptable specificity (0.558, 95% CI: 0.413-0.695). These findings support the complementary value of integrating proteomic and immunological biomarkers with clinical assessment to improve identification of patients. In the secondary analysis focused on patients with prolonged remission, model performance further improved, particularly with respect to specificity and overall accuracy.

**Table 2.** Predictive performance of clinical, biomarker, and combined models in the full intervention arm and in patients with prolonged remission.

| Model | Threshold | Accuracy<br>(95% CI) | Sensitivity<br>(95% CI) | Specificity<br>(95% CI) | PPV<br>(95%CI) | NPV<br>(95% CI) |
| --- | --- | --- | --- | --- | --- | --- |
| **Intervention arm** |  |  |  |  |  |  |
| Clinical | 0.4871 | 0.649<br>(0.546-0.744) | 0.644<br>(0.488-0.781) | 0.654<br>(0.509-0.780) | 0.680<br>(0.533-0.805) | 0.617<br>(0.464-0.755) |
| Biomarkers | 0.4361 | 0.691<br>(0.589-0.781) | 0.822<br>(0.679-0.920) | 0.577<br>(0.432-0.713) | 0.789<br>(0.627-0.904) | 0.627<br>(0.491-0.750) |
| Combined | 0.337 | 0.701<br>(0.6-0.79) | 0.867<br>(0.732-0.949) | 0.558<br>(0.413-0.695) | 0.829<br>(0.664-0.934) | 0.629<br>(0.497-0.748) |
| **Patients with prolonged remission** |  |  |  |  |  |  |
| Clinical | 0.4871 | 0.703<br>(0.585-0.803) | 0.644<br>(0.488-0.781) | 0.793<br>(0.603-0.920) | 0.590<br>(0.421-0.744) | 0.829<br>(0.664-0.934) |
| Biomarkers | 0.2844 | 0.784<br>(0.673-0.871) | 0.867<br>(0.732-0.949) | 0.655<br>(0.457-0.821) | 0.76<br>(0.549-0.906) | 0.796<br>(0.657-0.898) |
| Combined | 0.4311 | 0.784<br>(0.673-0.871) | 0.8<br>(0.654-0.904) | 0.759<br>(0.565-0.897) | 0.71 (0.52-0.858) | 0.837<br>(0.693-0.932) |

CI, confidence interval; DAS-28-CRP, disease activity score-28 (based on CRP); PPV, positive predictive value; NPV, negative predictive value

To evaluate the potential clinical utility of these models, decision curve analysis was performed. Across a range of clinically relevant threshold probabilities (20–40%), the combined model consistently demonstrated a higher net benefit compared with both the clinical and biomarker-only models (Figure 4C). To facilitate clinical interpretation, net interventions avoided (NIA) were also estimated. At a threshold probability of 30%, the combined model avoided approximately 16 inappropriate tapering decisions per 100 patients, representing 15 additional avoided interventions compared with the clinical model alone. Together, these findings support the added clinical value of combining molecular biomarkers with disease activity measures to guide biologic tapering strategies in RA.

## DISCUSSION

Although bDMARDs have revolutionized RA management by enabling sustained remission, their use is associated with dose-dependent side effects, treatment burden, and high costs [13]. Therefore, some clinical guidelines suggest considering dose reduction in patients with prolonged remission and under certain conditions, although others recommend maintaining dose continuation rather than reduction, based on low certainty of the evidence [5]. In this context, the OPTIBIO study reported higher relapse rates among patients who tapered bDMARDs compared with those maintained on standard doses, although many remained in remission [8]. These findings, together with recent systematic review and meta-analysis data [20], suggest that treatment de-escalation may be suitable only for selected individuals, highlighting the need to identify patients most likely to benefit from tapering. However, clear criteria for patient selection are still lacking, which underscores the need for biomarkers to guide dose reduction strategies [13].

Our study aimed to identify protein biomarkers associated with flare risk during bDMARD tapering, to support the accurate selection of patients who might safely undergo dose reduction. In the MS-based discovery phase using 44 OPTIBIO samples, baseline VSIG4 levels differed between patients who remained in remission and those who later flared. In the full cohort (N=194), baseline VSIG4 levels were significantly higher in patients who experienced a flare compared with those who remained in remission, with this association being particularly evident within the intervention arm. Elevated baseline VSIG4 was associated with increased flare risk and a higher likelihood of flare over time supporting its role as a candidate biomarker in the context of treatment tapering.

VSIG4, a member of the B7 family, regulates T cells through a receptor not yet identified and plays a critical role in immune suppression [21,22]. Soluble VSIG4 (sVSIG4), released during macrophage activation, inhibits complement activation by binding C3b and blocking the alternative complement pathway [23]. This function has been therapeutically exploited in rodent models of RA, where sVSIG4 reduced disease severity and prevented bone damage [24], and VSIG4-CD59 fusion proteins improved outcomes in adjuvant-induced arthritis [25].

In humans, VSIG4 is expressed by tissue-resident regulatory macrophages in both healthy tissue and RA synovium during remission but it is reduced in refractory RA [18]. Certain VSIG4 genetic variants have also been linked to RA severity [26]. Increased sVSIG4 levels have been reported active lupus nephritis [27] and have been associated with short-term mortality in spontaneous bacterial peritonitis [28]. Although further functional studies are needed, our findings reveal for the first time an association between VSIG4 and RA flares, highlighting its potential as a biomarker to identify patients unlikely to benefit from bDMARD tapering.

In addition, MS revealed modestly higher levels of S100A8 and S100A9 in patients who flared compared with those who remained in remission within the intervention arm. The S100A8/S100A9 heterodimer (calprotectin) correlates strongly with RA activity compared with traditional acute-phase reactants, but its predictive value for flares remains inconsistent. In our ELISA validation of the full cohort, baseline calprotectin levels did not differ significantly between patients who flared and those who did not, suggesting a limited role in this specific clinical context.

Anti-cytokine autoantibodies (ACAAs) are increasingly recognized as modulators of the immune responses. They can influence cytokine function by reducing or enhancing signaling activity, or by altering their half-life in circulation [29–33]. ACAAs are found in both healthy individuals and in various pathological conditions, including autoimmune diseases [34,35]. In autoimmune disorders, ACAAs may serve as prognostic indicators, as shown for anti-IL-8 and anti-IL-1α in RA, and anti-IL-6 in systemic sclerosis [36–38]. In our study, anti-IFNγ levels were associated with an increased risk of flare in patients undergoing bDMARD tapering, and also with earlier flares over time, supporting their potential role as predictive biomarkers.

IFNγ, produced by lymphocytes and natural killer cells, plays a central role in both innate and adaptive immunity. Its relationship with RA is complex, as it exerts both proinflammatory and regulatory effects [39]. In early RA, IFNγ promotes inflammation via the JAK-STAT pathway, inducing MHC class II expression, driving M1 macrophage polarization, and stimulating antimicrobial responses [40]. Conversely, in chronic RA, IFN-γ may exert regulatory functions, such as inhibition of Th17 differentiation and modulation of T-cell activation [41,42]. Furthermore, IFNγ can inhibit osteoclast formation by promoting degradation of TRAF6, an adapter protein in the RANKL pathway, thereby protecting against bone erosion in chronic inflammation [39,43]. Although anti-IFNγ autoantibodies have not been previously described in RA, they have been associated with immune dysregulation in other contexts [44–46]. Our findings support further investigation into their biological role in RA flares.

Among all clinical variables analyzed, only higher DAS28-CRP was independently associated with increased flare risk in the tapering arm of the study. When integrated into predictive models, VSIG4 and anti-IFNγ improved the performance of clinical prediction. Furthermore, the combined model showed better discrimination than the clinical variables alone and achieved a significantly higher sensitivity, which is a clear benefit given the clinical objective of minimizing inappropriate treatment tapering in patients at risk of flare. These results suggest that biomarker-informed models may enable more precise risk stratification and support personalized treatment optimization in RA.

Importantly, the added value of biomarkers was not limited to statistical performance but extended to clinical utility. Decision curve analysis demonstrated that the combined model provided a higher net benefit across clinically relevant threshold probabilities compared with the clinical model, indicating improved decision-making for treatment tapering. This translated into a reduction in inappropriate tapering decisions, with the combined model avoiding additional unnecessary interventions.

Although the primary analysis was conducted in the entire intervention arm, a secondary analysis focusing on patients with prolonged remission, representing those most likely to benefit from treatment optimization. This analysis showed improved discriminative performance, with AUC values reaching 0.839, suggesting that the proposed biomarker-based models may be particularly effective in identifying patients with stable long-term remission, who are the most suitable candidates for treatment de-escalation.

This study has some limitations. The main limitation is the lack of external validation in an independent cohort, owing to current unavailability of comparable datasets. Nevertheless, prospective validation in an external population is planned to confirm the reproducibility and generalizability of these findings. Larger studies will be needed to further refine biomarker performance and assess clinical applicability in broader patient populations. Despite these limitations, the study has important strengths. It is based on samples from a randomized clinical trial with a prospectively calculated sample size, ensuring methodological rigor and statistical robustness. The combination of discovery proteomics and targeted immunoassays provides a comprehensive and reliable approach for biomarker identification and validation. Most importantly, these results represent a step forward toward personalized medicine in RA, supporting safer and more effective bDMARD tapering strategies.

## CONCLUSIONS

In conclusion, this study identifies VSIG4 and anti-IFNγ autoantibodies as novel circulating biomarkers associated with RA flare during bDMARD tapering. Their combination with DAS28-CRP substantially improves predictive performance and provides clinically relevant benefit in decision-making, supporting the development of biomarker-guided strategies for personalized treatment optimization.

## LIST OF ABBREVIATIONS

ACAAs: Anti-cytokine autoantibodies
ACPA: Anti-citrullinated protein antibodies
ACR: American College of Rheumatology
AUC: Area under the curve
bDMARDs: Biological disease-modifying antirheumatic drugs
BMI: Body mass index
CI: Confidence interval
CRP: C-reactive protein
CV: Coefficient of variation
DAS28: Disease Activity Score in 28 joints
DAS28-CRP: Disease Activity Score in 28 joints based on C-reactive protein
DDA: Data-dependent acquisition
DIA: Data-independent acquisition
ELISA: Enzyme-linked immunosorbent assay
EULAR: European Alliance of Associations for Rheumatology
FDR: False discovery rate
HR: Hazard ratio
IFNγ: Interferon gamma
IgG: Immunoglobulin G
MFI: Median fluorescence intensity
NPV: Negative predictive value
OR: Odds ratio
PASEF: Parallel accumulation serial fragmentation
PPV: Positive predictive value
PRIDE: PRoteomics IDEntifications Database
RA: Rheumatoid arthritis
RF: Rheumatoid factor
ROC: Receiver operating characteristic
SDAI: Simplified Disease Activity Index
STRING: Search Tool for the Retrieval of Interacting Genes/Proteins
TNF: Tumor necrosis factor
VSIG4: V-set immunoglobulin-domain-containing protein 4

## DECLARATIONS

### Ethics approval and consent to participate

The Biologic Optimization Study (OPTIBIO) was a Phase IV, randomized, controlled, open-label, non-inferiority trial conducted in patients with RA in sustained remission on biological therapy (EudraCT 2012-004482-40). The study adhered to ethical principles in biomedical research and the current legislation in Spain, following Good Clinical Practice (ICH Topic E6, 2011), the Declaration of Helsinki, and the requirements of RD 223/2004. The Autonomous Ethics Committee of Galicia (CEIC) approved the study, and all patients provided written informed consent.

### Consent for publication

Not applicable.

### Availability of data and materials

The mass spectrometry proteomics data have been deposited to the ProteomeXchange Consortium via the PRIDE [14] partner repository with the dataset identifier PXD073940. Other datasets used and/or analysed during the current study are available from the corresponding author on reasonable request.

### Authors’ contributions

FJB and CRR conceived and designed the project. PQ, VC, PFP, RPG, LG, BA, NO, RR, and LL performed the research. PDG, VBB and DN analyzed the data. FJB, LL and CRR wrote the manuscript. All authors approved the final manuscript.

## Acknowledgements

Part of the protein analysis was performed by the ICTS “NANBIOSIS”, specifically by the Proteomics Unit of the CIBER-BBN at SERGAS (Spain).

