## Supplementary material for "Integrated serum proteomics and autoantibody analyses reveal a biomarker signature predictive of flare during biologic tapering in rheumatoid arthritis"

**Supplementary Table 1.** Baseline characteristics of patients included in the discovery phase conducted by MS. Provided as an Excel table.

**Supplementary Table 2.** List of proteins identified in the discovery phase by MS. Provided as an Excel table.

**Supplementary Table 3.** Results from the quantitative proteomic analysis in the control group. Provided as an Excel table.

**Supplementary Table 4.** Results from the quantitative proteomic analysis in the intervention group. Provided as an Excel table.

**Supplementary Table 5.** Univariate regression analysis of clinical variables in the intervention group.

| Variable | OR (95% CI) | p value |
|---|---|---|
| Age | 1.031 (0.999-1.065) | 0.061 |
| Sex | 1.013 (0.327-3.078) | 0.981 |
| BMI | 1.084 (1.009-1.172) | **0.033** |
| RF | 0.998 (0.413-2.431) | 0.996 |
| ACPA | 1.196 (0.504-2.881) | 0.686 |
| CRP | 5.586 (1.069-36.949) | 0.053 |
| DAS28-CRP | 4.444 (1.707-13.166) | **0.004** |

**Supplementary Table 6.** Logistic regression models and discriminative performance of clinical, biomarker, and combined models in the intervention arm and prolonged remission analysis.

| Model | Variable | OR (95% CI) | p | AUC (95% CI) | Adjusted AUC (95% CI) |
|---|---|---|---|---|---|
| **Intervention arm** | | | | | |
| Clinical | DAS28-CRP | 4.444 (1.707-13.166) | 0.004 | 0.666 (0.557-0.774) | 0.666 (0.566-0.785) |
| Biomarkers | VSGI4 | 4.03 (1.707-9.913) | 0.002 | 0.722 (0.62-0.823) | 0.713 (0.62-0.816) |
| | anti-IFNɣ | 1.757 (1.756-1.758) | 0.06 | | |
| Combined | VSGI4 | 2.992 (1.197-7.655) | 0.02 | 0.76 (0.666-0.854) | 0.738 (0.653-0.832) |
| | anti-IFNɣ | 1.809 (1.808-1.811) | 0.051 | | |
| | DAS28-CRP | 3.155 (1.122-9.904) | 0.037 | | |
| **Prolonged remission** | | | | | |
| Clinical | DAS28-CRP | 7.983 (2.276-37.138) | 0.003 | 0.725 (0.607-0.842) | 0.722 (0.61-0.849) |
| Biomarkers | VSGI4 | 4.597 (1.602-14.343) | 0.006 | 0.769 (0.656-0.883) | 0.758 (0.652-0.866) |
| | anti-IFNɣ | 3.835 (3.831-3.839) | 0.011 | | |

| | | | | | |
|---|---|---|---|---|---|
| Combined | VSGI4 | 3.975 (1.257-13.781) | 0.022 | 0.839 (0.747-0.931) | 0.815 (0.735-0.91) |
| | anti-IFNɣ | 5.067 (5.061-5.073) | 0.006 | | |
| | DAS28-CRP | 10.508 (2.385-65.875) | 0.005 | | |

ORs for anti-IFNγ were calculated per 1000 MFI units. Values are shown with 95% confidence intervals (CI). Bootstrap-adjusted AUC values were obtained using internal bootstrap validation.

**Supplementary Figure 1**

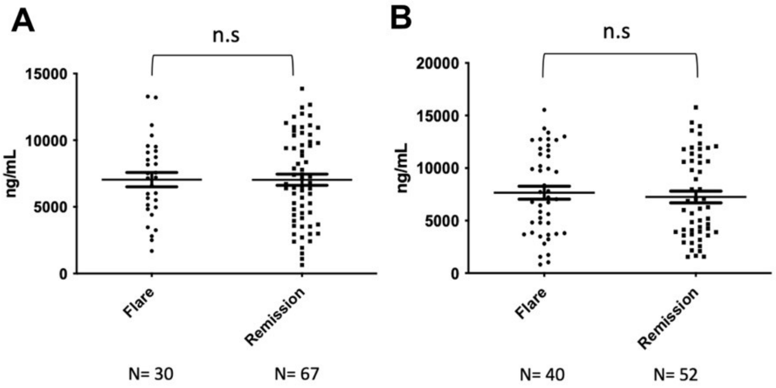


**Supplementary Figure 1. Baseline serum calprotectin (S100A8/S100A9) levels are not associated with flare during follow-up.** Calprotectin concentrations (ng/mL) measured at baseline are shown for (A) control arm patients, and (B) intervention arm patients, stratified by flare status. P values were calculated using Mann-Whitney U tests. Bars represent median and interquartile range. n.s., not significant.

**Supplementary Figure 2**

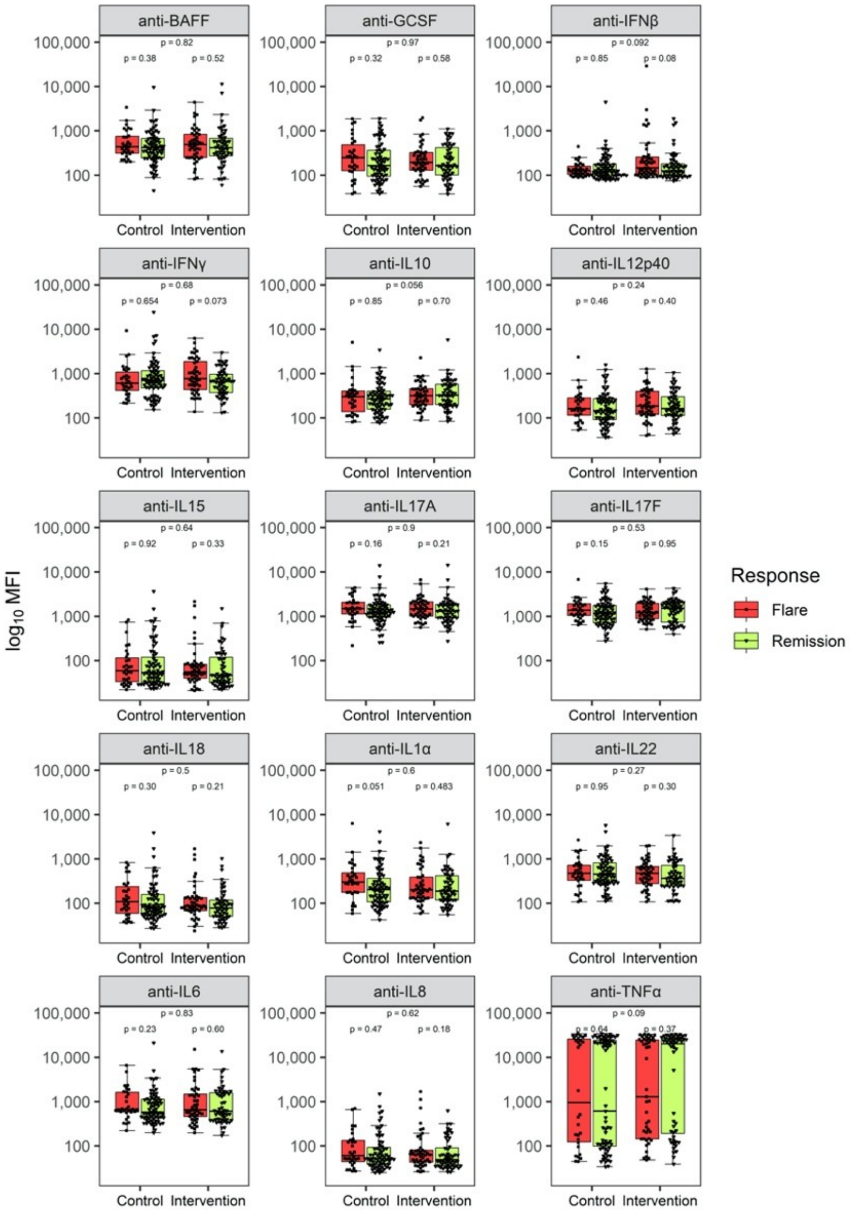


**Supplementary Figure 2. Baseline serum levels of anti-cytokine autoantibodies in the OPTIBIO cohort.** Boxplots comparing the levels of IgG levels against 15 cytokines (BAFF, G-CSF, IFNβ, IFNγ, IL-1α, IL-6, IL-8, IL-10, IL-12p40, IL-15, IL-17A, IL-17F, IL-18, IL-22, and TNFα) between flare and remission patients stratified by study arm (control vs. intervention). IgG levels are expressed as log10-transformed median fluorescence intensity (MFI) and P-values were calculated using Mann-Whitney U tests.
